# mslp: a comprehensive pipeline in predicting cancer mutations specific synthetic lethal partners

**DOI:** 10.1101/2021.12.15.472776

**Authors:** Chunxuan Shao

**Author notes:** **Corresponding author** Correspondence to Chunxuan Shao,.

## Abstract

**Background:** Mutation specific synthetic lethal partners (SLPs) offer significant insights in identifying novel targets and designing personalized treatments in cancer studies. Large scale genetic screens in cell lines and model organisms provide crucial resources for mining SLPs, yet those experiments are expensive and might be difficult to set up. Various computational methods have been proposed to predict the potential SLPs from different perspectives. However, those efforts are hampered by the low signal-to-noise ratio in simple correlation based approaches, or incomplete reliable training sets in supervised approaches.

**Results:** Here we present mslp, a comprehensive pipeline to identify potential SLPs via integrating genomic and transcriptomic datasets from both patient tumours and cancer cell lines. Leveraging cuttingedges algorithms, we identify a broad spectrum of primary SLPs for mutations presented in patient tumours. Further, for mutations detected in cell lines, we develop the idea of consensus SLPs which are also identified as screen hits, and show consistency impact on cell viability. Applied in real datasets, we successfully identified known synthetic lethal gene pairs. Remarkably, genetic screen results suggested that consensus SLPs have a significant impact on cell viability compared to common hits.

**Conclusions:** Mslp is a powerful and flexible pipeline to identify potential SLPs in a cancer context-specific manner, which might aid in drug developments and precise medicines in cancer treatments. The pipeline is implemented in R and freely available in github.

## Background

Synthetic lethality (SL) is defined as the reduced cell viability by the simultaneous inactivation of two genes, while perturbations of either of them only has no or mild impact. Since its first description in *Drosophila melanogaster*, this concept has gained much attention in cancer studies where one gene involved in the synthetic lethality relationship is mutated in patients [1]. The BRCA1/2-PARP1 in breast cancer might be the most famous and well studied synthetic lethality pairs, and recently ENO1-ENO2 has been reported in glioblastoma where ENO1 is deleted as a passenger gene in 1p36 locus [2, 3]. By targeting mutation related SLPs, it might be possible to inhibit growth of tumour cells in a genetic specific manner, where the normal cells surrounding tumour will be largely unaffected. Therefore, SLPs provide exciting insights on designing personalized treatments for individual patients based on the mutation profiles [4].

Large efforts have been made to identify putative SLPs related to mutations. Genome scale genetic screens have been extensively applied to both model organisms (e.g., yeast) and common cancer cell lines, adapting the shRNA knockdown or more recent Crispr-Cas9 knockout system [5]. Meanwhile, various computational methods have been proposed to predict SLPs from different perspectives and utilize a broad type of datasets, including metabolic, genomic and transcriptomic profiles in both cell lines and patients. DAISY provides a comprehensive approach to predict candidate SLPs integrating multiple sources, yet the selecting threshold based on Spearman correlation is not an optimal solution due to the nonlinear interaction among genes [6]. In addition, there is considerable discrepancy of mutations between cell lines and patients, *e*.*g*., some mutations in human patients are not detected in available cell lines. Based on the assumption that SLPs of mutations are amplified more frequently or deleted less frequently, MiSL is a novel approach to directly identify SLPs from patients genomic datasets, however, ignoring the scenario of co-expression between mutation and SLPs limits the scope of potential SLPs [7]. On the other hand, supervised machine learning techniques are popular choices in mining the synthetic lethal relations, but the performance of predictions suffers from the incomplete set of known SL datasets which are used as training data [8].

Here we present a comprehensive computational pipeline, mslp (Mutation specific Synthetic Lethal Partners), to identify consensus SLPs of mutations in a cancer context-specific manner through explicitly integrating genomic and transcriptomic data from patients and cell line genetic screen data. Firstly, we infer the primary SLPs based on two simple yet complementary assumptions 1) gene expression of mutations correlate with SLPs in wide type patients, 2) up-regulated expression of SLPs compensate for the loss of function of the mutation [6, 7]. Two computational modules, correlationSLP and compensationSLP, are derived from state-of-art statistical methods to realize the assumptions with the ever-increasing patients omics data. Further, in spite of complex gene-gene interaction, we hypothesize that genetic perturbations targeting mutation specific SLPs would likely reduce cell viability. Thus, for the recurrent mutations detected in both patient tumours and cancer cell lines in which genetic screen data are available, we developed the idea of consensus SLPs under two constraints, 1) primary SLPs are screen hits, 2) consistent impact on cell lines with the targeted mutations. When applied to real datasets, we found that knock-down of predicted consensus SLPs has a significant impact on cell viability. Thus, mslp provides a novel approach to study mutation specific SLPs with great flexibility.

### Implementation

Mslp is implemented in R and could be run parallelly (Fig. 1a). The major steps are 1) collect and prepare input data; 2) identify the primary SLPs in compensationSLP and correlationSLP modules from patients tumor datasets; 3) identify consensus SLP for mutations detected in both patient tumours and cell lines. The following sections cover details in each step, in addition, a thorough tutorial with example datasets from breast invasive carcinoma is provided in the package vignette.

**Figure 1:**
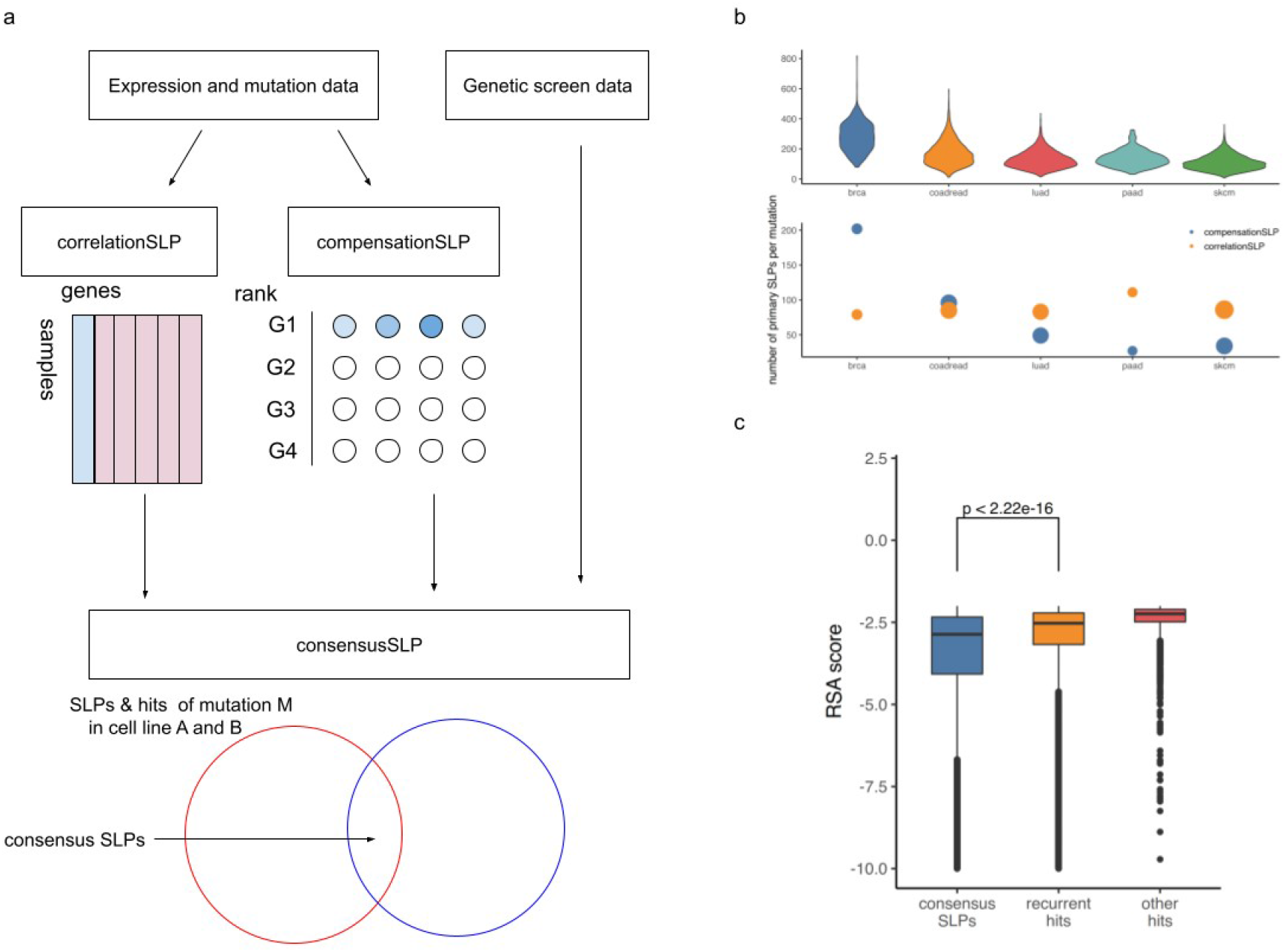
Pipeline overview and results in real datasets. **a** A schematic flowchart of mslp pipeline. Primary SLPs identified by correlationSLP and compensationSLP are combined together to obtain consensus SLPs. **b** Upper panel, distribution of primary SLPs per mutation among cancers; lower panel, primary SLPs per mutation identified by each module, the size of bubble represents number of mutations in cancers. **c** Boxplot of RSA value of hits in different categories. *P*-value is calculated by Student’s t-test.

### Data preparation

Four types of data would be necessary to run a complete analysis of mslp: expression (including normalized value and z-score value which reflects relative expression) and mutation profiles of patients tumours in interested cancer types, as well as mutation and genetic screen datasets in corresponding cell lines. Mslp provides utilities to preprocess data downloaded from cBioPortal, including steps of genes/hypermutations filtering and mutations selection [9]. Cancer cell line profiles could be downloaded from various databases like Cancer Cell Line Encyclopedia (CCLE) and Project DRIVE [10, 11].

### Identify primary SLPs

In the compensationSLP module, we employ the Rank Products algorithm (RP) to identify up-regulated genes that might compensate for the loss of function of mutations, utilizing the mutation and z-score profiles in patients tumour datasets. RP is a non-parametric and robust method of identifying differentially regulated genes. For each mutation *M* detected in two or more patients, we build a z-score by patients matrix which only includes patients with *M*. To reduce the false positive rate, genes that are co-mutated with *M*, or are expressed with positive z-score in less than 20% of patients are removed from downstream analysis. In addition, we filter out genes with higher z-score in wide type patients than in patients with *M*. The preprocessed matrix is used as the input to the “RankProducts” function in the R package RankProd using the option “calculateProduct = FALSE”, which is able to analyze large datasets and more robust to extreme values [12, 13]. The *P*-value of up-regulated genes are adjusted for multiple comparisons by the “Benjamini and Hochberg” method.

In the correlationSLP module, we utilize GENIE3 to identify genes that highly correlate with mutations in wide type patients. Briefly, GENIE3 performs random forest based regression to select the most likely regulations between predictor and response variables, based on variable importance [14]. For each *M* used as the response variable, we focus on genes positively correlated with *M* (Pearson correlation coefficient *r* > 0) in wild type patients as predictors. Candidate SLPs are selected with variable importance value >= 0.001 as the primary results. We derive a receiver operating characteristic (ROC) based approach on repetition runs of GENIE3 to assess the significance of the genes. Shortly, GENIE3 is applied on a subset of randomly selected mutations repeatedly. The top ranked SLPs are selected via the RP method with adjusted *P*-value <= 0.001 for each mutation, and labeled as “TRUE” SLPs. We then perform the ROC analysis on candidate SLPs in each repetition and calculate the Youden index, where SLPs are split into “TRUE” and “FALSE” groups (“TRUE” genes are defined above) [15]. We define the threshold as the average Youden index value across genes and repetitions.

#### Identify consensus SLPs

Large scale genetic screen (*e*.*g*., Project DRIVE) and cell line mutation profiles (*e*.*g*., CCLE) enable us to examine the reductions of cell viability upon perturbations of predicted SLPs for mutations detected in both patients and cell lines. For each *M* detected in two or more cell lines, we select the overlapping of primary SLPs and screen hits, and then calculate the Cohen’s Kappa coefficient to determine the consistency of those genes pair-wisely among cell lines. The consensus SLPs are selected based on thresholds on Kappa coefficient and/or *P*-value.

## Results

We applied mslp to five cancer types with most abundant cell line profiles in CCLE, including breast invasive carcinoma (BRCA), colorectal adenocarcinoma (COADREAD), lung adenocarcinoma (LUAD), pancreatic adenocarcinoma (PAAD) and skin cutaneous melanoma (SKCM). The patients datasets (“data_RNA_Seq_v2_expression_median.txt”, “data_RNA_Seq_v2_mRNA_median_Zscores.txt”, “data_mutations_extended.txt” and “data_CNA.txt”) were downloaded from cBioPortal [9], genetic screen data were downloaded from Project DRIVE. Interestingly, we found that the importance thresholds used in the correlationSLP module were comparable among the cancer datasets (Supplementary Fig. 1). We identified 165 primary SLPs on average for each mutation (*P*-value <= 0.01 in compensationSLP module, importance >= 0.0016 in correlationSLP module) (Fig. 1b). About 0.2% SLPs are identified by both compensationSLP and correlationSLP modules, confirming our pipeline could identify a broad spectrum of candidates. To test whether those SLPs are truly context-specific in each cancer, we calculated the average percentage of overlapping SLPs for mutations detected in at least two cancer cohorts pair-wisely. The percentage is highly skewed towards zero, with a mean value of 11%, confirming the specificity of our approach (Supplementary Fig. 2). Remarkably, we successfully identify the well-known synthetica lethal pairs of BRCA1/2-PARP1 in BRCA.

Next we tried to identify consensus SLPsfor mutations detected in both patients and cell lines. RSA values provided by Project DRIVE indicate the statistical significance of whether the target plays a role in cell viability, and in our analysis hits were selected with RSA <= -2. On average 33 cell lines were used in the downstream analysis among cancer types. 11 consensus SLPs are identified per mutation (Kappa coefficient >= 0.6 and adjusted *P*-value <= 0.1). To examine the impact of consensus SLPs on cell viability, we divided hits into three groups, consensus SLPs, recurrent hits that are shared by two or more cell lines excluding consensus SLPs, and other hits. Remarkably, consensus SLPs are found to have significantly smaller RSA compared to recurrent hits, suggesting that mslp could identify the most vulnerable targets (Fig. 1c).

## Conclusion

We provide an end-to-end pipeline to identify putative SLPs related to specific tumour mutations, explicitly integrating both genomic and transcriptomic data in patients and genetic screen data in related cancer cell lines. Based on two simple assumptions, we employ cutting-edge algorithms to predict the primary SLPs with a novel approach to control the false positive rate. Further, for mutations detected in both human tumours and cancer cell lines, we identify consensus SLPs which are screen hits and persistent among cell lines. Applied to several real tumour datasets, we showed that the consensus SLPs had a significant impact on cell viability compared to recurrent hits. With ever-growing expression and genetic screen datasets, mslp might aid in hypothesis testings in precise medicines and drug developments.

## Availability

https://github.com/ccshao/mslp

## Acknowledgements

The author thanks Dr. Hai-kun Liu for helpful discussions, Dr. Tobias Schmelzle for providing the RSA score data.

## Author information Affiliations

Division of Molecular Neurogenetics, German Cancer Research Center (DKFZ), DKFZ-ZMBH Alliance, Im Neuenheimer Feld 280, Heidelberg 69120, Germany

## Contributions

CS designed the project, implemented it with R codes and wrote the manuscript.

## Corresponding author

Correspondence to Chunxuan Shao.

## Ethics declarations

## Ethics approval and consent to participate

Not applicable.

## Consent for publication

Not applicable.

## Reference

1. Bridges CB. The Origin of Variations in Sexual and Sex-Limited Characters. Am Nat. 1922;56:51–63.

2. Lord CJ, Ashworth A. PARP inhibitors: Synthetic lethality in the clinic. Science. 2017;355:1152–8.

3. Muller FL, Colla S, Aquilanti E, Manzo VE, Genovese G, Lee J, et al. Passenger deletions generate therapeutic vulnerabilities in cancer. Nature. 2012;488:337–42.

4. Gupta A, Ahmad A, Dar AI, Khan R. Synthetic Lethality: From Research to Precision Cancer Nanomedicine. Curr Cancer Drug Targets. 2018;18:337–46.

5. Nijman SMB. Synthetic lethality: General principles, utility and detection using genetic screens in human cells. FEBS Lett. 2011;585:1–6.

6. Jerby-Arnon L, Pfetzer N, Waldman YY, McGarry L, James D, Shanks E, et al. Predicting Cancer-Specific Vulnerability via Data-Driven Detection of Synthetic Lethality. Cell. 2014;158:1199–209.

7. Sinha S, Thomas D, Chan S, Gao Y, Brunen D, Torabi D, et al. Systematic discovery of mutation-specific synthetic lethals by mining pan-cancer human primary tumor data. Nat Commun. 2017;8:ncomms15580.

8. Das S, Deng X, Camphausen K, Shankavaram U. DiscoverSL: an R package for multi-omic data driven prediction of synthetic lethality in cancers. Bioinformatics. 2019;35:701–2.

9. Cerami E, Gao J, Dogrusoz U, Gross BE, Sumer SO, Aksoy BA, et al. The cBio Cancer Genomics Portal: An Open Platform for Exploring Multidimensional Cancer Genomics Data. Cancer Discov. 2012;2:401–4.

10. McDonald ER, de Weck A, Schlabach MR, Billy E, Mavrakis KJ, Hoffman GR, et al. Project DRIVE: A Compendium of Cancer Dependencies and Synthetic Lethal Relationships Uncovered by Large-Scale, Deep RNAi Screening. Cell. 2017;170:577-592.e10.

11. Barretina J, Caponigro G, Stransky N, Venkatesan K, Margolin AA, Kim S, et al. The Cancer Cell Line Encyclopedia enables predictive modelling of anticancer drug sensitivity. Nature. 2012;483:603.

12. Hong F, Breitling R, McEntee CW, Wittner BS, Nemhauser JL, Chory J. RankProd: a bioconductor package for detecting differentially expressed genes in meta-analysis. Bioinformatics. 2006;22:2825–7.

13. R Core Team. R: A Language and Environment for Statistical Computing. Vienna, Austria: R Foundation for Statistical Computing; 2020.

14. Huynh-Thu VA, Irrthum A, Wehenkel L, Geurts P. Inferring Regulatory Networks from Expression Data Using Tree-Based Methods. PLoS ONE. 2010;5:e12776.

15. Robin X, Turck N, Hainard A, Tiberti N, Lisacek F, Sanchez J-C, et al. pROC: an open-source package for R and S+ to analyze and compare ROC curves. BMC Bioinformatics. 2011;12:77.

