## Supplemental File for "mslp: a comprehensive pipeline in predicting cancer mutations specific synthetic lethal partners"

Chunxuan Shao

Division of Molecular Neurogenetics, German Cancer Research Center (DKFZ), DKFZ-ZMBH Alliance, Im Neuenheimer Feld 280, Heidelberg 69120, Germany

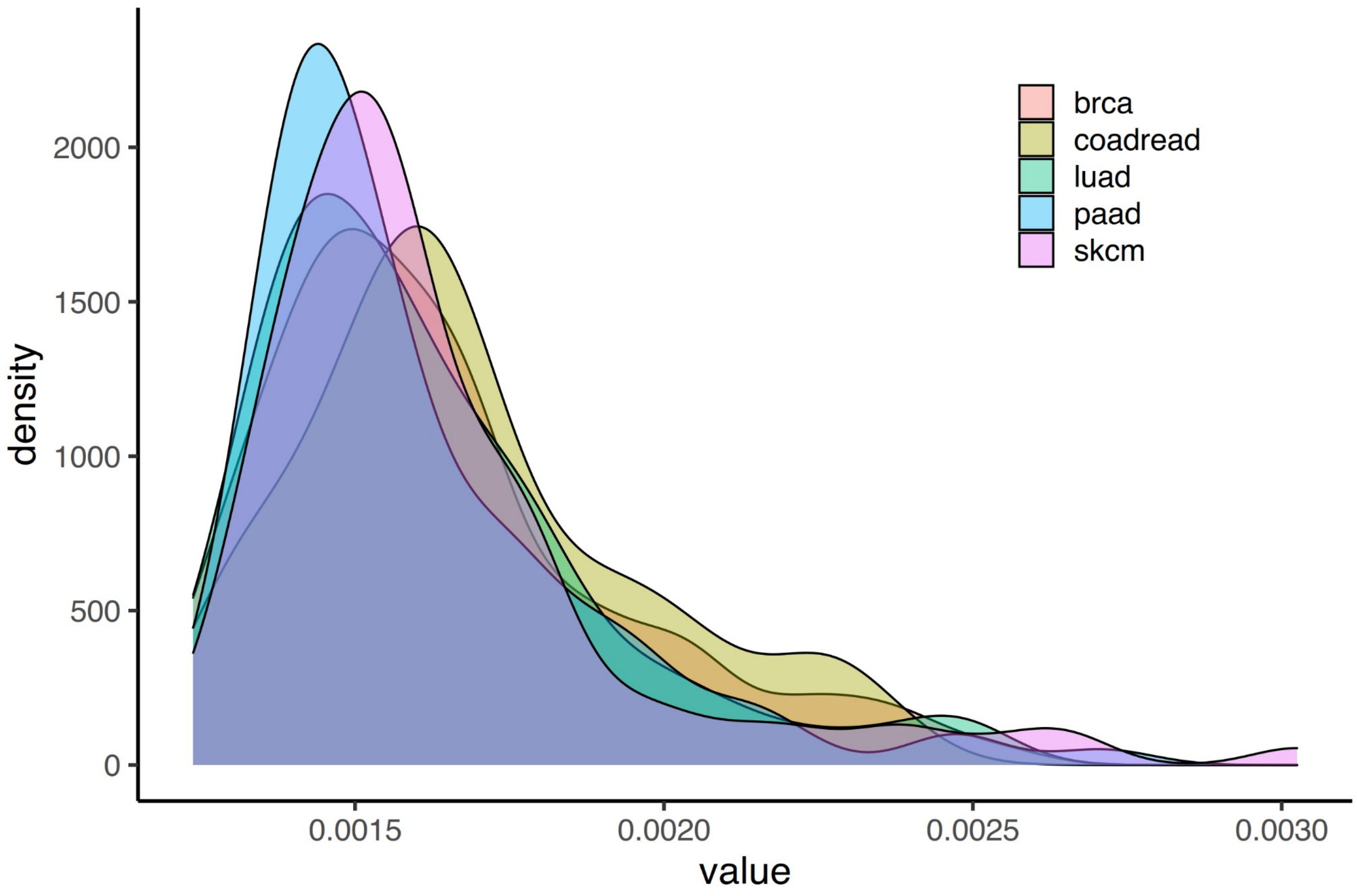


**Supplementary Figure 1.** Estimated importance threshold. The distribution of importance threshold estimated in five different cancers by the ROC approach.

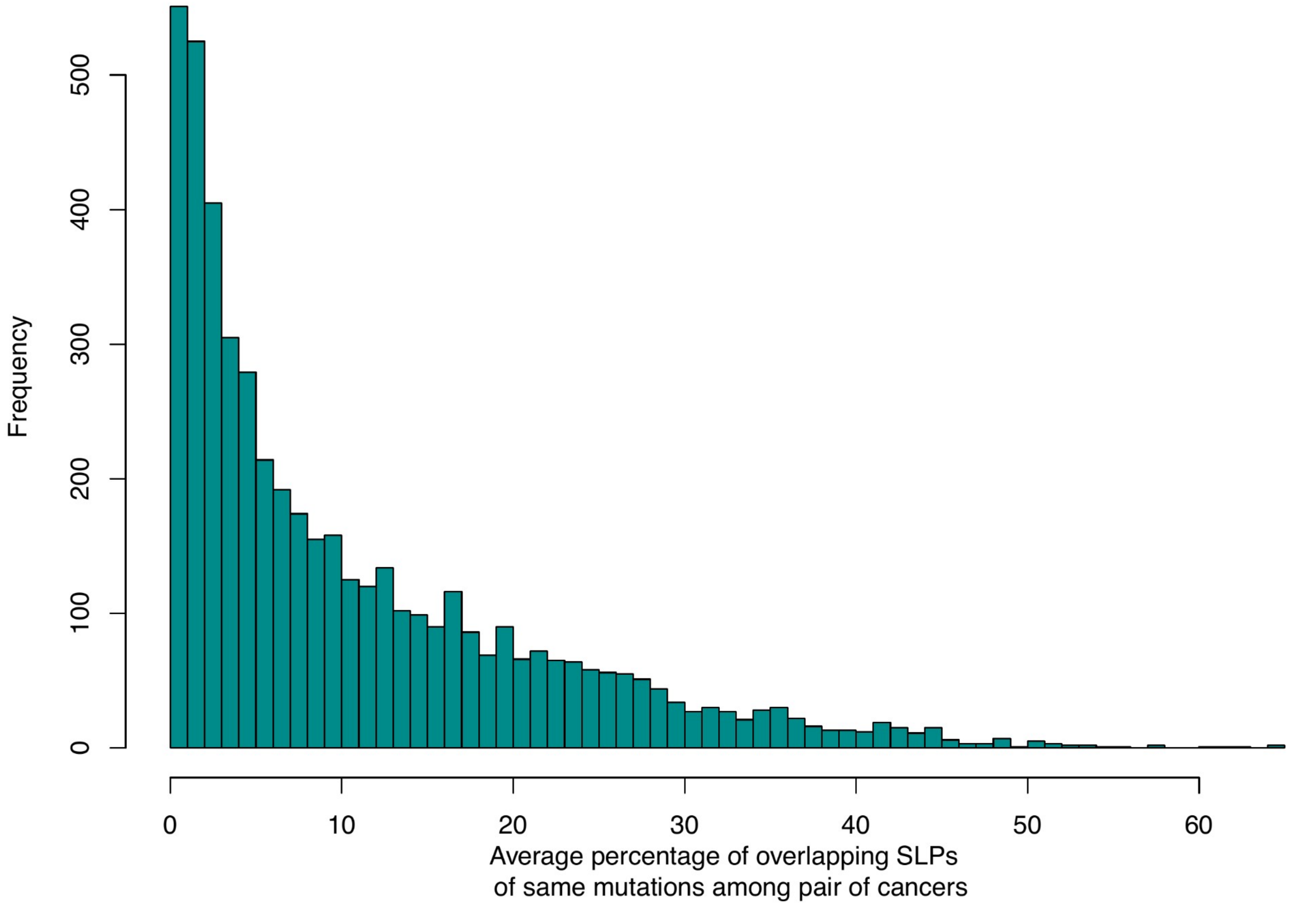


**Supplementary Figure 2.** Histogram of percentage of overlapping SLPs. For each mutation found in two or more cancer types, we calculated the percentage of common primary SLPs against the average number of primary SLPs.
